# DNA polymerase α-primase can function as a translesion DNA polymerase

**DOI:** 10.1101/2025.07.02.662785

**Authors:** Ryan Mayle, Roxana Georgescu, Michael E. O’Donnell

**Affiliations:** HHMI and The Rockefeller University, New York, New York 10065

**Keywords:** DNA polymerase alpha, Primase, DNA translesion bypass, DNA replication, DNA repair, Replication fork

## Abstract

Replication of cellular chromosomes requires a primase to generate short RNA primers to initiate genomic replication. While bacterial and archaeal primase generate short RNA primers, the eukaryotic primase, Polα-primase, contains both RNA primase and DNA polymerase (Pol) subunits that function together to form a >20 base hybrid RNA-DNA primer. Interestingly, the DNA Pol1 subunit of Polα lacks a 3’-5’ proofreading exonuclease, contrary to the high fidelity normally associated with DNA replication. However, Polδ and Polδ synthesize the majority of the eukaryotic genome and both contain 3’-5’ exonuclease activity for high fidelity. None the less, even the small amount of DNA produced by Pol1 in each of the many RNA/DNA primers during chromosome replication adds up to tens of millions of nucleotides in a human genome. Thus it has been a longstanding question why Pol1 lacks a proofreading exonuclease. We show here that Polα is uniquely capable of traversing common oxidized or hydrolyzed template nucleotides and propose that Polα evolved to bypass these common template lesions when they are encountered during chromosome replication.

**Significance statement:** Eukaryotic Polα-primase contains DNA polymerase (Pol1) and RNA primase subunits that together synthesize a >20 nucleotide hybrid RNA-DNA primer. Bacteria and archaea only require a dozen or less RNA residues to prime DNA synthesis. Therefore, why do eukaryotes require the additional DNA? We propose, and demonstrate here that Pol1, which lacks a proofreading 3’-5’ exonuclease, is capable of traversing some common template lesions produced in the normal hydrolytic and metabolic oxidative environment of cells. Thus, we hypothesize that Pol1 activity within the eukaryotic primase evolved to help replisomes bypass template damage. Bypassed damaged sites can be dealt with by repair processes after replication has occurred.

## Introduction

Pol⍺-primase (referred to here as Pol⍺) is a 4-subunit complex common to all eukaryotes. The largest subunit, Pol1, is the DNA polymerase; it lacks a 3’-5’ exonuclease (1). The Pri1 and Pri2 subunits synthesize an 8-10 nucleotide RNA primer that is extended by Pol1 by 10-20 nucleotides of DNA, yielding a hybrid RNA-DNA primer of around 25 nucleotides (2). The Pol12 subunit acts as a scaffold that organizes the other subunits in formation of the RNA-DNA primer (3, 4).

Pol⍺ was the first polymerase known to function in chromosomal synthesis, and when it was discovered to contain intrinsic RNA priming activity Pol⍺ was believed to perform both leading and lagging strand synthesis. Later studies identified Polδ and Polε, which are now known to perform the majority of lagging and leading strand synthesis, respectively (5, 6). Thus, the current view is that Pol⍺ is the primase that generates a hybrid RNA-DNA primer to initiate DNA synthesis of the leading strand and to initiate the numerous lagging strand fragments (7, 8).

Short RNA primers are used to initiate DNA synthesis in bacteria and archaea (9). It remains a mystery why Polα uses RNA and DNA synthesis to make hybrid RNA-DNA primers. Adding to this mystery is the question of why Pol1 lacks a 3’-5’ nuclease, needed for high fidelity, especially given that the DNA made by Polα constitutes about 1.5% of the yeast genome. This translates to tens of millions of bases in humans (10). Furthermore, Pol1 is essential and one documented role outside of priming bulk replication is at telomeres (11). Polα might have evolved to have a DNA polymerase for use during telomeric synthesis. Almost all bacteria and archaea utilize circular chromosomes, possibly avoiding need for Pol1 (12). However, it is also possible that Pol1 of Polα is important for some other role during replication of long eukaryotic chromosomes, as explained below.

A possible role for eukaryotic Polα‘s Pol1 polymerase is suggested by its shared feature of lacking a proofreader, as is the case for many translesion synthesis (TLS) DNA polymerases (13, 14). These include Polθ (A family) (15), Polζ (16) (B family), Polβ (17) (X family) and all Y family TLS polymerases, including bacterial Pols IV and V, and eukaryotic Polη, Polκ and Rev1 (18) (19). The absence of a 3’-5’ exonuclease may conceivably help TLS Pols proceed over template lesions because they can either stall or go forward but cannot excise backward (13, 20). Thus it seems reasonable to propose that the absence of a 3’-5’ exonuclease in Polα, an enzyme required for replisome progression, may facilitate TLS of some DNA lesions that the replisome frequently encounters, like the frequent oxidative and hydrolytic damage that would occur in all cells of an animal. This could be a first line mechanism for damage tolerance, while the primary TLS enzymes, namely Polζ, Polη, and Rev1 in yeast, would be the predominant means of lesion bypass.

Ability to bypass template lesions is expected to result in a lower fidelity of DNA synthesis, but the ability to bypass frequent base damage would enable the replisome to continue progression and the lesions could be corrected by post replication repair. To explore its potential TLS function, we assayed Polα for ability to bypass some commonly produced lesions: alkylated thymine glycol, an abasic site, and the oxidative damaged 8-oxoG, and compared the results with use of the replicative high-fidelity Polδ and Polδ. As a control we utilized the less common cyclobutane thymine dimer (CPD) that is known from genetic studies to be bypassed in yeast by Polη or Polζ.

We find here Polα does indeed contain TLS activity for some frequently expected DNA lesions. In fact, a side-by-side comparison shows similar efficiencies of Polα with TLS Polζ for bypass of thymine glycol and abasic sites. In this regard, it is important to note that Polα functions in the context of the replisome and therefore it likely encounters lesions at replication forks before other TLS Pols. For example, thymine glycol, abasic sites and 8- oxoG lesions can be repaired by pathways outside of encounter by a replisome, but sometimes a replisome might encounter these frequent lesions before they are repaired. 8-oxoG is error prone because it can pair either correctly with C, or incorrectly with A, and thus is sometimes thought of as a mutagenic form of damage requiring post-replication repair rather than being a replication blocking lesion, or bypassed by enzymes like Polη in a relatively error free manor (21, 22). Bypass of a CPD lesion is also known to utilize Polη (23). However, a CPD lesion can be bypassed by either Polη or by Polζ in yeast (16, 24), stimulated by PCNA (proliferating cell nuclear antigen) (25). In this report, we examine Polα for ability to bypass thymine glycol, abasic, 8-oxoG and CPD template lesions.

In sum, this report presents a study of a continuum of lesion severity to Polα’s bypass ability. While we find that Polα can bypass frequent lesions in DNA, consistent with lack of a 3’-5’ exonuclease, it is blocked by other lesions such as a CPD. The report also documents an unexpected PCNA independent, but RFC (Replication Factor C) dependent process of thymine glycol lesion bypass by Polδ, revealing an additional function of the clamp loader besides loading of PCNA onto DNA. In overview, we suggest that Polα should be added to the list of TLS DNA polymerases.

## Results

### Both Polδ and Polδ can utilize an RNA primer

We first examine a very simple reason that could explain why Pol⍺ generates an RNA-DNA primer. Either Polδ and/or Polε, the two major eukaryotic replicative Pols, may not be able to use a 3’ RNA terminus as efficiently as a DNA 3’ terminus. We tested this by comparing extension using RNA primed and DNA primed substrates (Fig. 1A). The results show no detectable difference in extension of a DNA vs RNA primed template by either Polδ or Polε (Fig. 1B-C). Therefore, it seems unlikely that Pol1 evolved to generate a hybrid RNA/DNA primer specifically to provide a DNA terminus for use by either Polε or Polδ.

**Figure 1:**
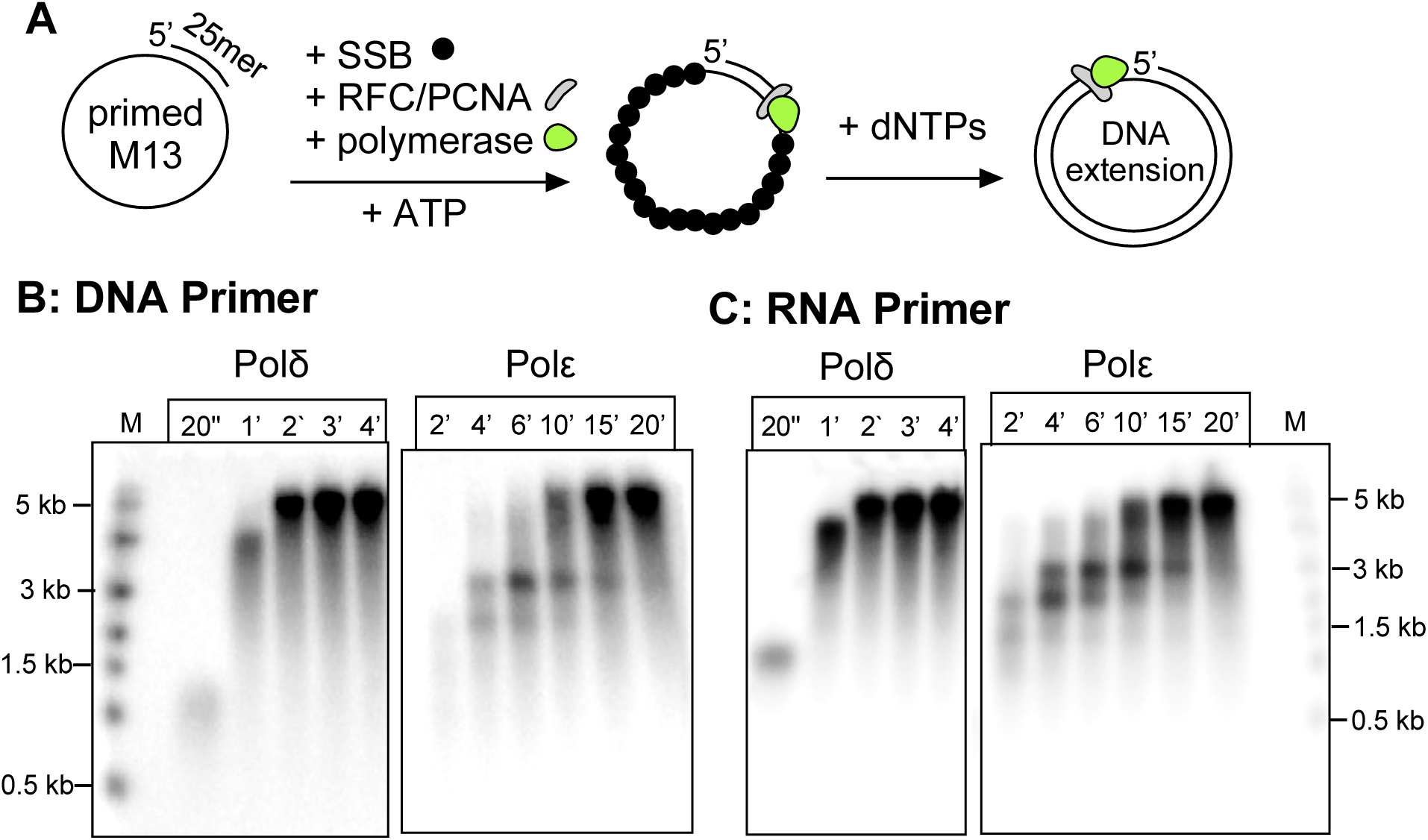
**Comparison of RNA vs DNA primers for extension by Pols δ and ε**. A) Reaction scheme, PCNA was incubated with RFC for loading of PCNA onto DNA or RNA primed M13mp18 ssDNA coated with SSB along with either Polδ or Polε, followed by addition of dNTPs for DNA synthesis. B) Time course of Polδ and Polε extension of DNA primed M13 mp18 ssDNA. C) Time course of Polδ and Polε extension of RNA primed M13 mp18 ssDNA. Products were separated by denaturing agarose gel electrophoresis.

### Bypass of a thymine glycol lesion

A recent biochemical study showed that a template thymine glycol (Tg), an oxidized form of thymine that occurs quite frequently *in vivo,* can be bypassed by Polδ-PCNA (i.e. using Polδ, RFC and PCNA) (26). However, the study did not address the possibility that Pol⍺ might bypass this lesion. It remained possible that the intrinsic tolerance for Tg at a replisome could, at least in part, be provided by the exo deficient Pol⍺.

To explore Pol⍺’s ability to bypass a template Tg lesion, we designed a primer extension assay where the template strand contains a Tg that must be bypassed to synthesize full length products (Fig. 2A). Details for this substrate, as well for all others used in the is study, are shown in Figure S1. The unextended primer, full length extension product, and intermediate stalled at the Tg lesion can be clearly resolved using denaturing polyacrylamide gel electrophoresis. End labeled oligos corresponding to these three forms were run as reference/control lanes to compare to extension products (Fig. S2).

**Figure 2.**
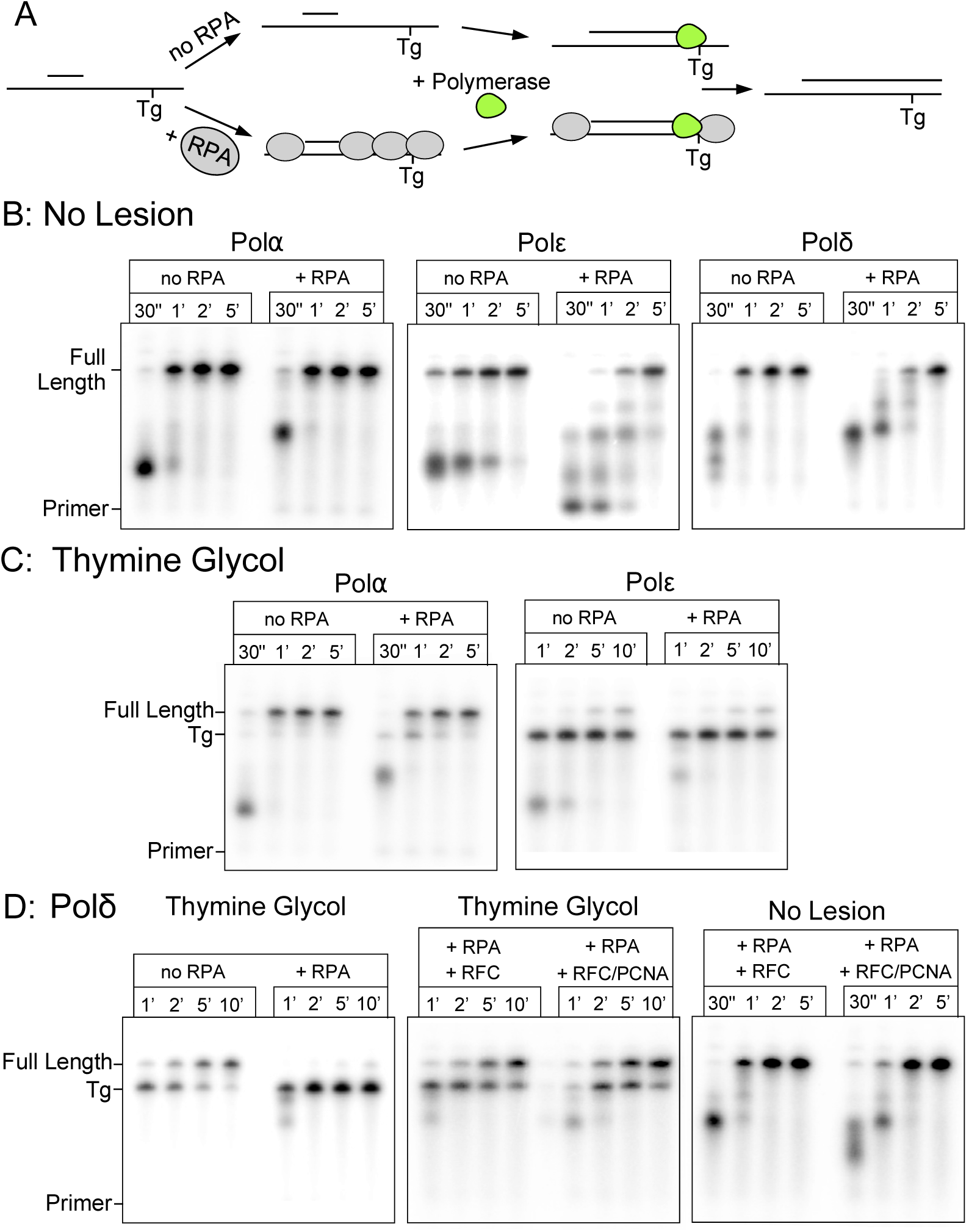
Polα efficiently bypasses a template thymine glycol lesion. A) Example reaction scheme for primer extension assay. B-D) Primer extension products separated by denaturing PAGE containing the indicated polymerase and DNA substrate. B) No lesion control. C) Thymine glycol lesion. D) Polδ analysis with RFC (minus PCNA) or RFC+PCNA and the indicated substrates.

Using comparable templates with an identical sequence, but with thymine rather than thymine glycol (Tg), shows that Pol⍺ proceeds on this substrate with no stall products at the size expected for Tg blocked extension (Fig. 2B). This is an important observation to be confident that extension intermediates of that size are in fact stalled at the lesion, rather than the possibility of unrelated sequence specific pausing. Extension assays on the Tg substrate reveal that Pol⍺ is capable of TLS over a template Tg lesion. As shown in (Fig. 2C), Polε only minimally bypasses the template lesion, compared to rapid and efficient bypass by Pol⍺ with no requirement for additional cofactors. Pol⍺ bypass is still efficient when RPA (Replication Protein A) is present, though marginally slowed. This stands in contrast to Polδ, where bypass of Tg is observed on naked DNA but is strongly inhibited by RPA. This result is consistent with a recent study which showed that Polδ can bypass Tg in the presence of RFC-PCNA (26).

Interestingly, the earlier study of Polδ/RFC/PCNA bypass of a Tg lesion (26) used a linear blunt end synthetic primed template, and thus PCNA might have been loaded onto DNA but then escaped over the blunt end of the DNA, leaving some ambiguity to the results, and leaving open the possibility that either RFC or PCNA might be the only protein responsible. Therefore we assayed Polδ and Polε in the presence and absence of RFC/PCNA. Our Tg bypass substrate includes sufficient ssDNA on either side of the primer for RPA binding to ensure loaded PCNA is retained on the DNA.

### RFC alone, without PCNA, stimulates Tg bypass by Polδ

We observe RFC-PCNA stimulated Polδ bypass of Tg (e.g. when RPA is present), but also unexpectedly saw that RFC alone is sufficient for bypass in the absence of PCNA (Fig. 2D). Comparable assays with the “no lesion” substrate demonstrate that RFC can, by itself, stimulate Polδ extension independent of PCNA (Fig. 2D). That RFC, independent of PCNA, stimulates Polδ has not previously been demonstrated, and these new findings suggest there is a functional interaction between RFC and Polδ. This expands the possible roles of clamp loaders beyond loading of the replicative PCNA clamp. In contrast to Polδ, Polε is not positively impacted by RFC-PCNA and still shows only weak Tg bypass (Fig. S3).

### Bypass of a template abasic site

Loss of a nucleotide base is a frequent lesion caused by hydrolysis (27, 28). Therefore the next lesion we tested for TLS activity by the replicative Pol⍺ was an abasic site. A substrate containing a template strand abasic site, similar to that used to analyze the Tg substrate, was compared versus use of the substrate containing no abasic site. These DNA substrates are shorter than that used for the Tg lesion (Fig S1, Table S1), and control reactions that lack any lesion confirmed this primed template is completed with minimal pausing within 1 minute of initiation (Fig. S2).

Interestingly, only Pol⍺ could bypass an abasic site (Fig. 3A), while Polδ and Polε could not. Significantly, Polδ could not do so even in the presence of RFC-PCNA, and Polε showed no ability to bypass an abasic site under any condition tested (Fig. 3A, Fig. S3). While Pol⍺ bypass is slowed by RPA, there is substantial full length product formed, which is not true for Polδ or Polε even in the absence of RPA.

**Figure 3:**
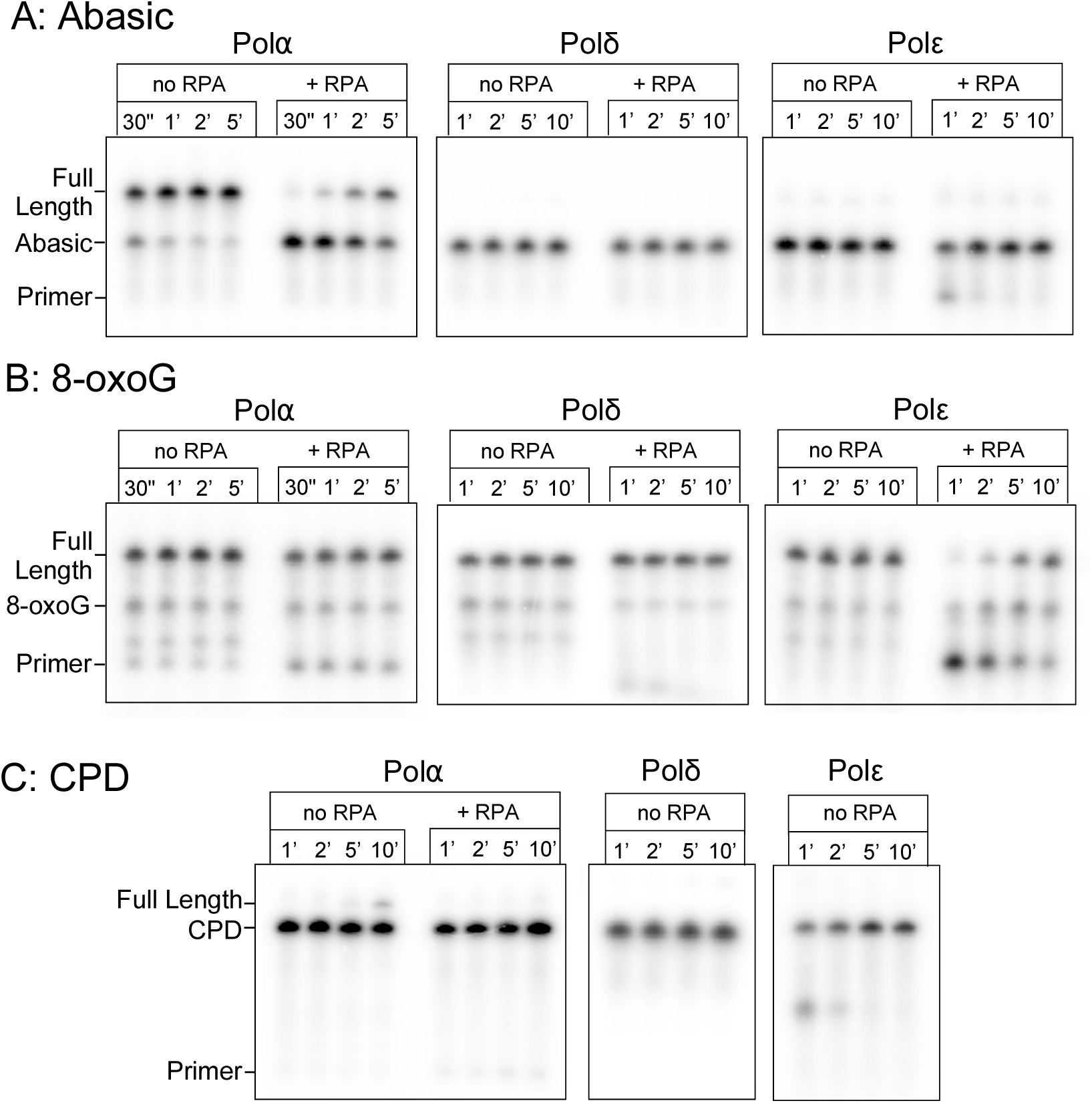
Polα can bypass an abasic site. Primer extension on template strands carrying an abasic site using either Pol⍺, Polδ or Polε, in the presence and absence of RPA (as indicated above the gel). Positions of markers are shown next to the gel. (A), CPD (B), or 8-oxoG (C). Products generated by the indicated polymerase were separated by denaturing PAGE.

### Bypass of a template 8-oxoG lesion

8-oxoG, an oxidized guanine base, is a mutagenic base that readily mispairs with adenine. The expectation is that a template 8-oxoG can be bypassed but results in errors due to its base pairing ambiguity. We observe that all three replicative polymerases, Pol⍺, Polδ and Polε, synthesize past this lesion with approximately the same efficiency (Fig. 3B). While bypass appears comparable for all pols on naked DNA, addition of RPA specifically inhibits Polε. This difference in efficiency may have relevance *in vivo*, but broadly, 8-oxoG is not a strict block to any of these polymerases, consistent with previous studies of both yeast and bacterial polymerases (29, 30).

### A cyclobutane pyrimidine dimer (CPD) blocks each of the replicative Pols

Ability of Pol⍺ to bypass Tg, abasic sites and 8-oxoG template lesions demonstrates that Pol⍺ is distinct from Polδ and Polε in bypass activity for some common template lesions that are expected in all cells of an organism. One or both of these template lesions pose a block to Polδ and Polε, but not Pol⍺, indicating that Pol⍺ could serve a useful role during bulk replication upon replisome encounter with these common lesions.

Cyclobutane pyrimidine dimers (CPD) are among the most common forms of UV induced DNA damage and have been shown to require both Polη and Polζ for translesion bypass in the human system, or by either Polη or Polζ in yeast (18). Indeed, for CPD lesions formed outside the replication cycle, the multiprotein nucleotide excision repair pathway has been characterized for removing these lesions. This is a form of damage known to inhibit replication progression, and therefore the expectation is that none of the replicative polymerases alone will be capable of efficient CPD bypass.

Indeed, we observe that Pols ⍺, δ, and ε alone are incapable of efficient CPD bypass. We do observe detectable, although inefficient, bypass of a template CPD by Pol⍺ (Fig. 3C). The presence of RFC and PCNA do not provide CPD bypass ability for either Polδ or Polε (Fig. S3). Bypass of a CPD lesion by Pol⍺ is notably slower and less efficient than bypass of Tg and abasic lesions. But given sufficient time it is clearly occurring, while Polδ and Polε show no detectable bypass. It is important to note however that RPA strongly inhibits CPD bypass by Pol⍺. Therefore, it remains to be seen whether bypass of a CPD lesion by Pol⍺ has physiological relevance. Nonetheless, these observations expand the scope of what Pol⍺ is intrinsically capable of.

### Rate of Pol⍺ TLS bypass compared to a known TLS Pol

This report shows that Pol⍺ displays TLS bypass activity, well above the activity by the Polδ and Polε replicative enzymes. To contextualize the relative proficiency of Pol⍺ in lesion bypass, we compared its activity to Polζ, a robust TLS enzyme in yeast. To test bypass of the Tg lesion, we primed the template just before the lesion to eliminate the time-lag effects of upstream fill-in on the extension time courses. Pol⍺ is able to bypass the Tg lesion in a similar time frame to Polζ, with both completing bypass and extension within 20-30 seconds (Fig. 4A).

**Figure 4:**
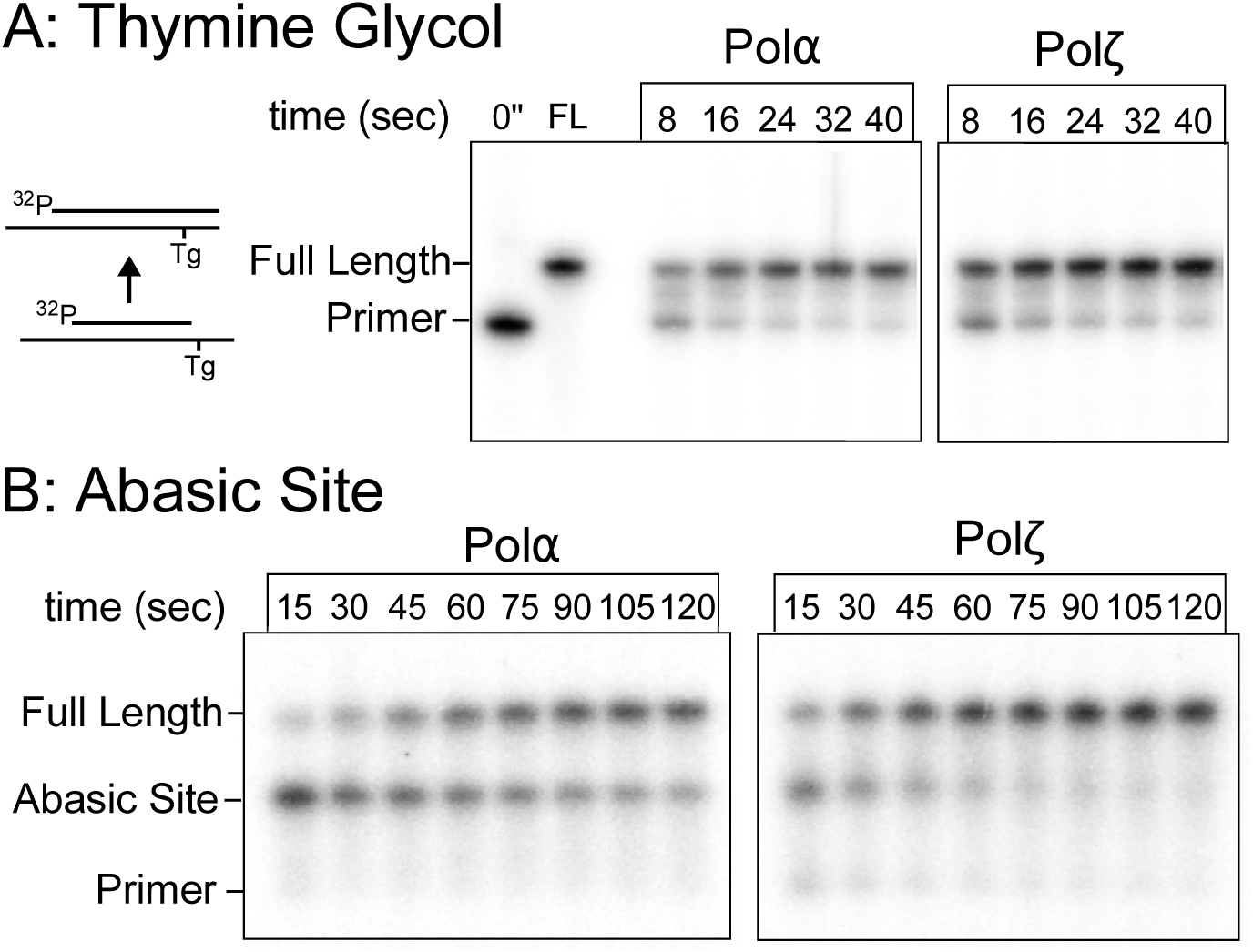
**Pol⍺ bypasses Tg and abasic sites at a comparable rate to Polζ**. Primer extension time-courses catalyzed by Pol⍺ or Polζ on: A) an abasic site primed DNA, and B) a Tg lesion substrate. Extension products were separated by denaturing PAGE.

Comparison of Pol⍺ and Polζ for TLS bypass at an abasic site indicates that Pol⍺ takes 2-4 times longer to perform bypass compared to Polζ points (Fig 4B), indicating that incorporation at the abasic site is slower compared to extension on undamaged DNA. Both enzymes have an easier time bypassing Tg compared to an abasic site, with minimal pausing at a Tg site but more prominent pausing at the abasic site, particularly for Pol⍺.

These results show that at certain commonly expected lesions, Pol⍺ possesses inherent TLS activity roughly comparable to an established TLS polymerase and therefore we propose that Pol⍺ could contribute to DNA damage tolerance during replication.

## Discussion

### Why do eukaryotes use a >20 nucleotide RNA-DNA primer?

As shown here, both eukaryotic replicative DNA polymerases, Polδ and Polε, are equally capable to utilize an RNA or DNA primer. This implies that use of 3’ RNA vs 3’ DNA is not the reason that Pol⍺-primase has evolved the ability to make an RNA-DNA hybrid primer. While one may question whether the >20 nucleotide length of the RNA-DNA primer is required, we have been unable to test this directly because 7-8 RNA oligos (i.e. the length of Pri1/Pri2 products) are not stably annealed to the DNA. However, we note that the primase of bacteriophage T4, which utilizes a sliding clamp and clamp loader only generates a 5 nucleotide RNA primer (stabilized by T4 primase-helicase) (31). Moreover, the *E. coli* primase makes an RNA primer of approximately a dozen ribonucleotides, which is sufficient for clamp/clamp loader function (1, 7, 9). Interestingly, archaea utilize a complex that is homologous to Pri1/2 of eukaryotes, but lacks a Pol1 or Pol12 homologue (32). Indeed, the data of Fig. 1 confirm that PCNA loading by RFC is not compromised by use of an RNA versus a DNA primed site. Therefore we propose that Pol⍺ has evolved the Pol1 activity for extending a primed site in eukaryotes for some specialized purpose(s) in eukaryotic replication, beyond simply a site for loading PCNA.

### Why does eukaryotic Pol⍺-primase lack proofreading exo activity?

An unanswered question in the replication field is why Pol⍺’s Pol1 subunit, that lacks 3’- 5’ exonuclease proofreading activity, has been retained throughout evolution in eukaryotes. The Pol1 gene is essential and besides DNA synthesis, Pol1 is known to be involved in structural roles such as nucleosome transfer during replication (1). The essential nature of Pol1 prevents simple deletion and examination of Pol1 with respect to mutability, considering the mutant would not be viable. Thus, one would need to identify a separation of function mutant that prevents Pol1 catalytic activity while leaving the structural role(s) of Pol⍺-primase intact. Such a mutant would allow for studying what role Pol⍺’s TLS abilities may have, as well as further characterizing other repair roles that have been suggested for the Pol1 subunit of Pol⍺. This possibility is currently being tested. Regardless, an important, and potentially essential role for Polα is at telomeres, where it is important for proper replication of chromosome ends (11).

It is interesting to note that Pol⍺ lacks a proofreading exonuclease yet is documented to replicate 1.5 % (150,000 bases) of the eukaryotic S.c. yeast genome (10). Extrapolation to the length of a human cell genome brings this extension number to over 40 Mb. The lack of a proofreader in Pol⍺, combined with its significant contribution to genomic DNA synthesis, suggests some type of proofreading exists. While one might imagine this proofreading could be mismatch repair, it has been demonstrated that Polδ proofreads errors made by Pol⍺ (33).

In this report, we propose that a benefit of having DNA Pol1 within Pol⍺-primase that does not contain proofreading activity, is that lack of a 3’-5’ exonuclease may enable the replisome to bypass some common template lesions caused by metabolic oxidative and/or hydrolytic damage, such as formation of thymine glycol adducts and abasic sites. A possible TLS use for Polα could enable genomic replication to proceed in certain situations, rather than pausing and waiting for other TLS enzymes to be recruited at more difficult and unique lesions (e.g. bulky lesions and interstrand cross-links). This hypothetical scenario is illustrated in Figure 5.

**Figure 5.**
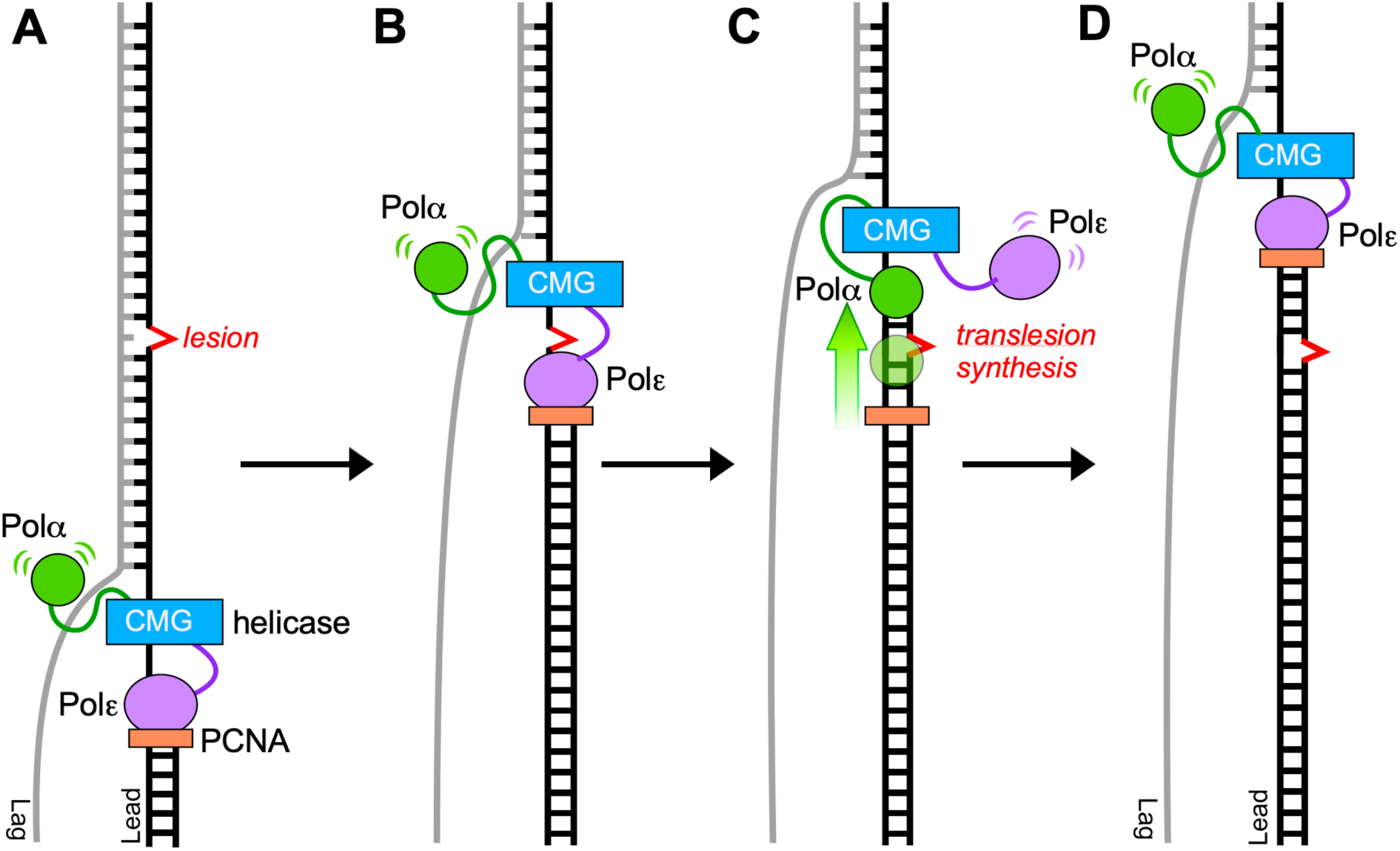
**Hypothetical use of Pol⍺ in lesion bypass on the leading strand**. While the CMG helicase may be expected to unwind DNA having a small lesion, the high fidelity replicative Pols are unlikely to bypass template lesions. The illustration focuses on the leading strand, although similar actions are presumed to occur on the lagging strand. Specifically, A) Pols ⍺ and ε bind simultaneously to the CMG-replisome components. B) Polε-PCNA stalls at a template lesion (e.g. such as a Tg or abasic site) and C) exchanges positions with Pol⍺; each Pol is observed to travel with the replisome via several protein- protein attachments (41), enabling lesion bypass. D) After bypass, Polε, documented to be strongly attached to CMG (42), remains bound to CMG and reassumes its position as the leading strand replicase after Pol⍺ bypass is complete. The lesion can be repaired later by post replication repair processes.

### Why does eukaryotic primase contain a DNA polymerase?

We can only speculate on the possible reason that eukaryotic cells utilize a Pol1 within Pol⍺ that lacks a proofreading nuclease. The archaea domain of life evolved on the same branch as current eukaryotes after the last universal common ancestor (LUCA) (34). Therefore it is interesting to note that archaea do not contain homologues of the Pol1/Pol12 subunits of eukaryotes, but they do contain homologues to Pri1/2 for primase activity (34). Archaea typically contain a circular chromosome(s), while eukaryotes typically have linear chromosomes. Hence it is tempting to speculate, in light of Pol⍺’s use at telomeres (11), that Pol⍺-primase contains the Pol1 DNA polymerase to maintain the telomeric ends of eukaryotic linear chromosomes, and might be dispensable for circular genomes. However, the actual role of Pol1 within Pol⍺ is not certain.

DNA damage tolerance is important to all cells, and this report indicates that the eukaryotic Pol⍺ contains an inherent DNA damage tolerance mechanism. This DNA damage tolerance may allow cells to proceed with replication after the replication fork encounters certain frequent DNA lesions due to hydrolysis and oxidative metabolism, as illustrated in Fig 5 on the leading strand as an example. Polζ and Polη are dedicated to the task of TLS bypass, but Pol⍺ lesion bypass may be more efficient for lesions that are encountered by ongoing replication forks. In fact Polδ-PCNA has been demonstrated to bypass Tg lesions (26), and we show here that Pol⍺ possesses TLS activity over Tg and other DNA lesions, even without PCNA. Pol1 is an essential gene, and thus it is not simple to obtain genetic evidence of its involvement in TLS. However, our results suggest Pol1 might be relevant to lesion bypass and therefore might shed light onto why this enzyme has evolved to be an integral part of the eukaryotic replisome.

Despite the observations noted in this report, we cannot be confident they explain why Pol⍺’s DNA Pol1 has been retained throughout evolution. In addition to its catalytic activity, Pol1 is an important structural gene within Pol⍺ that assists the transfer of nucleosomes from parental to daughter strands (35, 36). Given that Pol1 is an essential gene, it is not trivial to separate Pol1 catalytic functions from non-essential structural functions without first defining a separation of function Pol⍺ mutant that disrupts DNA Pol1 activity while leaving structural functions intact, including interaction with Pol12 and the RNA priming Pol1/2 subunits. A mutant such as this, if it were obtained, would enable the study regarding what role Polα’s synthetic and TLS activities may have that are separate from its roles in priming, nucleosome deposition, or other replisome contacts.

### Proposed evolution of Pol⍺

Related to the question of whether Pol1 activity is essential *in vivo* is the question of what role it may have had in the past. It could be that Pol⍺ was at some point the primary polymerase in eukaryotic cells, with additional DNA polymerases evolving later. Perhaps a unified polymerase-primase for replication was an efficient configuration for replicating both leading and lagging strands without need of other DNA polymerases or a dedicated primase. Evolution of the high fidelity and PCNA coupled DNA Polε and Polδ may have provided sufficiently higher fidelity compared to Pol⍺ and thereby utilized Pol⍺ as a primase and as a TLS polymerase that primes the fork, but also helps advance the fork over common lesions. Thus we propose Pol⍺ may be utilized for TLS bypass.

In summary, Pol⍺ may have been a versatile early polymerase capable of replicating the leading and lagging strands, and possibly also for TLS damage tolerance. However, Pol⍺ may have become relegated to a priming role when PCNA/RFC evolved for providing processive replication to other B family DNA polymerases (i.e. Pols δ and ε). If this were the case, Pol⍺ may no longer be required to fulfill as many roles as it once did for genomes early in evolution but may still carry the intrinsic biochemical activities to do so.

## Acknowledgements

We thank the National Institute of General Medical Sciences of the National Institutes of Health for support to M.E.O. (GM149862), the Howard Hughes Medical Institute (M.E.O.), and the Breast Cancer Research Institute (M.E.O.).

## Author contributions

M.E.O., R.M. and R.G. conceived the project, M.E.O., R.M., and R.G. designed research; R.M. and R.G. performed research; M.E.O., R.M., and R.G. analyzed the data; R.M. and M.E.O. wrote the manuscript.

## Materials and Methods

### Materials

^32^P-ATP was obtained from Perkin Elmer Life Sciences (Waltham, Massachusetts). Unlabeled dNTPs and rNTPs were purchased from GE Health-care. Concentration of protein samples were estimated by Bradford Protein stain (Bio-Rad Labs, Hercules, CA) using BSA as the standard. DNA Oligonucleotides were purchased from IDT (Integrated DNA Technologies, Coralville, IA). M13mp18 ssDNA and T4 polynucleotide kinase were purchased from New England Biolabs (Ipswich, MA).

### Proteins

All protein expression vectors in this report are deposited into Addgene and the corresponding Addgene numbers are noted below. *S. cerevisiae* PCNA was obtained by expression in *E. coli* as described (Addgene #239197) (37). The 3 subunit *S. cerevisiae* RPA was expressed in E. coli (Addgene #240742 and 240743) and purified as described (38).

*DNA Polymerases*: Purification of all the nuclear yeast DNA Pols, and their corresponding SDS PAGE analysis, were reported in our earlier study (39). We note here the Addgene accession information for the various expression plasmids we constructed for production of the clamp, the RPA and for the nuclear yeast DNA Pols α, δ, ε and Polζ. All genes for yeast expression were under control of the Gal1/10 promotor, enabling either 1 or 2 genes to be expressed by each plasmid. The Addgene numbers for the yeast integration vectors that express the four genes of Pol α-primase are: Pol1 #241975, Pol12 #241976, Pri1/2 #241977. Polα was induced and purified from cells as previously described (38). The 3- subunit Polδ was expressed in yeast using two plasmids (Addgene #’s for Polδ are: Pol3 #239198, Pol31/Pol32 #239199) and cell growth, induction and purification was as described (38). Purification of *S. cerevisiae* 4-subunit Polδ was from yeast expression of integrated genes and purified as described earlier (40).

The 4 subunit Pol ζ was also expressed by recombinant means in yeast, purified and analyzed by SDS PAGE as documented in our earlier report (39). Briefly, Pol ζ contains 4 subunits (Rev3/Rev7/Pol31/Pol32) and 3 plasmid vectors were integrated for expression: Rev3 containing a C-terminal 3X FLAG tag was cloned into pRS402/GAL (Ade^+^) (Addgene #241258). Rev7 was cloned into pRS405/GAL (Leu^+^) (Addgene #241259), and pRS403/GAL (His^+^)) for expression of the Pol31 and Pol32 subunits (Addgene #239199).

**DNA substrates:** Please refer to Table S1 for all DNA oligos used in this study. Primed template substrates for primer extension assays were constructed by annealing a 5’ ^32^P end labeled primer to a template strand. First, primers were 5’ end labeled using T4 polynucleotide kinase and γ-P^32^-ATP at a sub-stoichiometric ratio of ATP to DNA, leaving only a negligible amount of unincorporated ATP. Primed substrates were annealed by combining the two oligos at a ratio of 1.2:1 template strand:primer in TE + 100 mM NaCl (final concentration), heated to boiling, then allowed to cool slowly to room temperature. Oligos were paired as follows: No Lesion Substrate 1: Pr1 + LeadNL-long; No Lesion Substrate 2: Pr1 + LeadNL-short; Tg substrate 1: Pr1 + Lead-Tg; Tg substrate 2: Std1 + Lead-Tg; CPD substrate: Pr1 + Lead-CPD; Abasic substrate 1: Pr1 + Lead-Ab; Abasic substrate 2: Std3 + Lead-Ab; 8-oxoG substrate: Pr1 + Lead-8oxoG.

#### Primer Extension on Oligonucleotide Primed Substrates

The buffer for DNA synthesis assays was 20 mM Tris-Acetate pH 7.5, 4% glycerol, 0.1 mM EDTA, 40 μg/mL BSA, 5 mM DTT, 10 mM MgSO_4,_ 100 μM dNTPs and 20 nM polymerase (unless otherwise noted). When included, RPA was added to a final concentration of 30 nM. A DNA substrate mix containing all reagents except for DNA polymerase was first incubated for 5 minutes at 30℃ to allow RPA to bind. For consistency, this step was included across experiments, even for reactions that did not contain RPA. DNA synthesis was then initiated by adding the indicated DNA Pol. For reactions with RFC (5 nM final) and/or PCNA (20 nM final), ATP was present at 1 mM and RFC/PCNA were added with at the same time as polymerase. Reactions were allowed to proceed at 30℃ and timed aliquots were collected at the indicated times (typically 30”, 1’, 2’, 5’ or 1’, 2’, 5’, 10’) and quenched by addition of an equal volume of stop buffer (1% SDS, 50 mM EDTA, 8% glycerol, .01% bromophenol blue, 83% formamide). A variation of the extension assay with more rapid time points (to characterize the rate of bypass) used the same protocol, except reactions were at 27°C and the indicated DNA Pol was present at 10 nM.

To observe extension products, the quenched timed aliquots were boiled for 5 minutes and products were analyzed in 12% denaturing urea PAGE gels. Size standards in each gel include the starting substrate taken before addition of polymerase (0 time) to identify the position of unextended ^32^P-primer, a ^32^P-oligonucleotide corresponding to full length extension, and a ^32^P-oligonucleotide corresponding to extension to the lesion. These ^32^P- oligonucleotides are referred to as Std1 (to the lesion) and Std2 (full length) for the Tg, CPD, and No Lesion-long substrates, and are referred to as Std3 (to lesion) and Std4 (full length) for the shorter abasic, 8-oxoG, and No Lesion-short substrates. After electrophoresis was complete, gels were exposed to a phosphor imaging screen for at least 16 h then scanned with an Amersham Typhoon.

### Primed M13 Replication Assays

A total of 1 nM primed M13mp18/D25 or M13mp18 /R25, 0.8 µM SSB, 20 nM PCNA, 10 nm RFC and 20 nM Polδ or Polε, were incubated with 100 µM ATP for 5 min at 30°C. The reaction buffer contained 20 mM TrisCl (pH 7.5), 8 mM magnesium acetate, 50 mM KGlutamate, 10% glycerol, 30 µg/ml bovine serum albumin and 2 mM dithiothreitol. The replication reactions were started by the addition of 1 mM ATP, 60 µM each of dATP, dCTP, dTTP, 20 µM dGTP and 33 nM α-[^32^P] dGTP. The reactions were quenched with an equal volume of 40 mM Tris–HCl (pH 8.0), 0.2% SDS, 100 mM EDTA. For the analysis of the size of extension products, samples were subjected to alkaline agarose gel electrophoresis.

---

**Figure S1:**
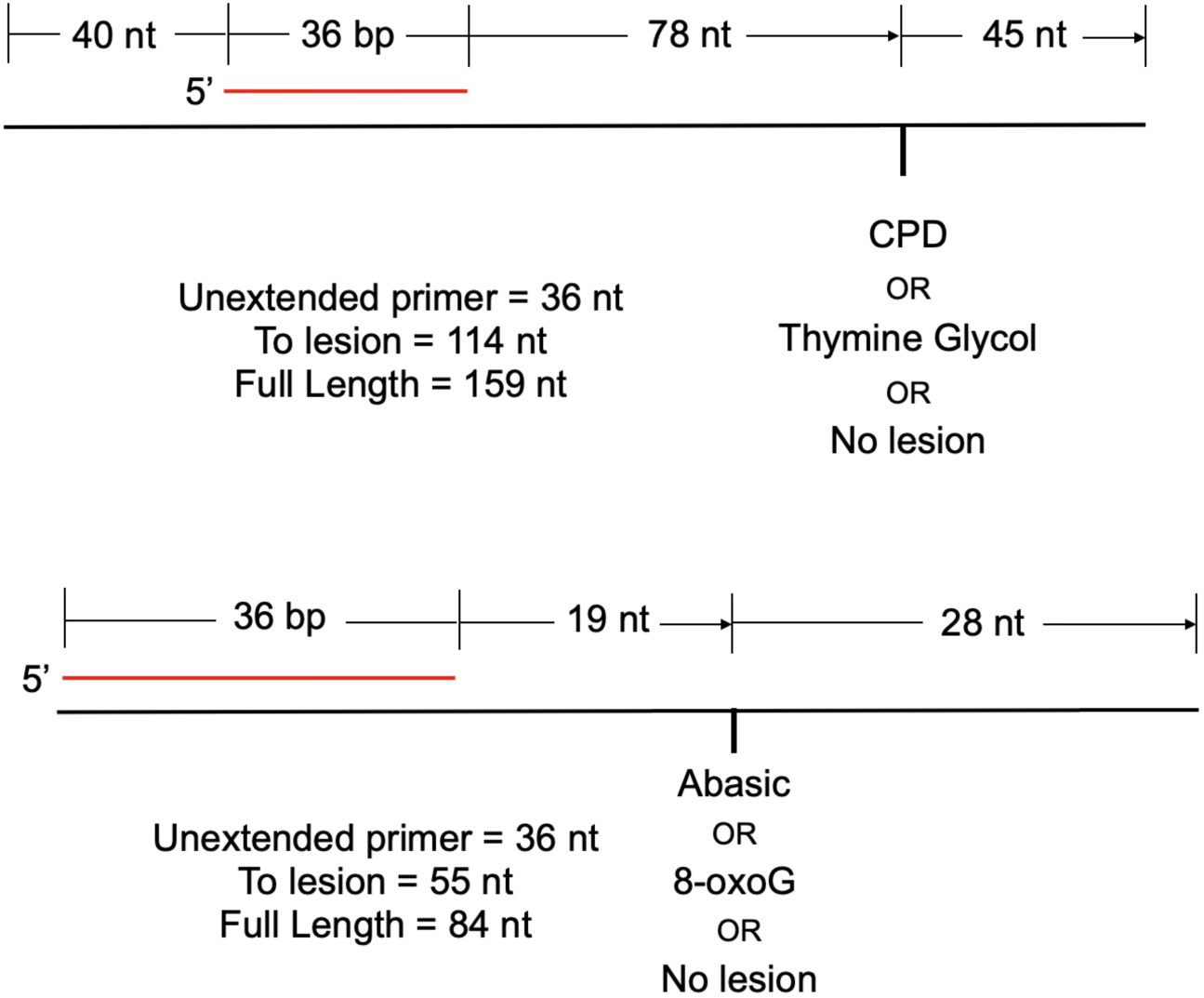
**Diagrams for primer extension substrates**. Details of the sizes for each section of the substrates used in our assay along with lengths of benchmark products. The primer, in red, is 5’ end labeled with ^32^P-ATP.

**Figure S2:**
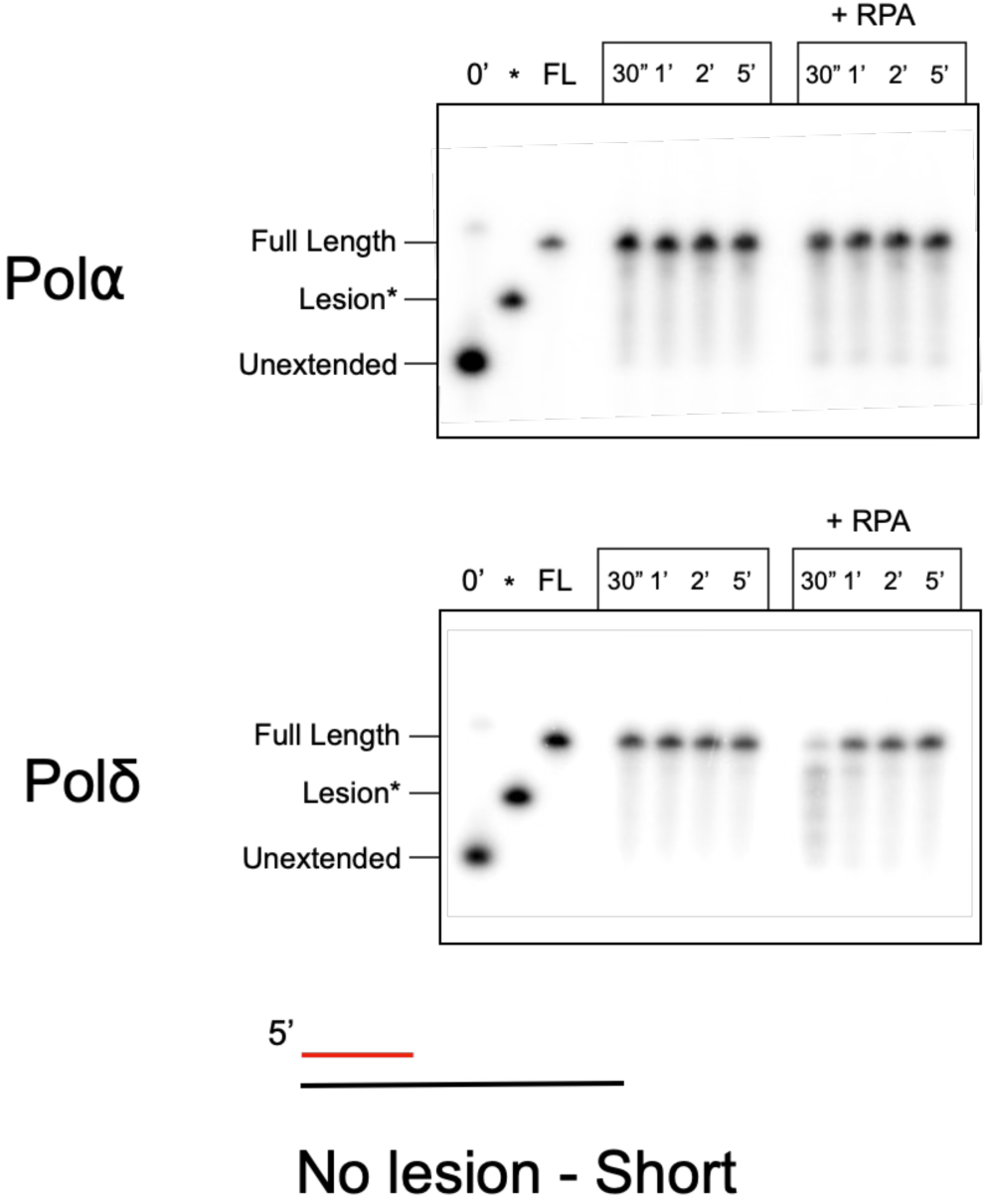
Primer extension assays on the short no lesion substrate. Extension products generated by Pol⍺ or Polδ with and without RPA. The * indicates products of a size corresponding to extension up to a lesion in the equivalent lesion substrates. This is included as a reference to show no pausing occurs at this location when no lesion is present. The ^32^P-ATP end-labeled primer is indicated in red.

**Figure S3:**
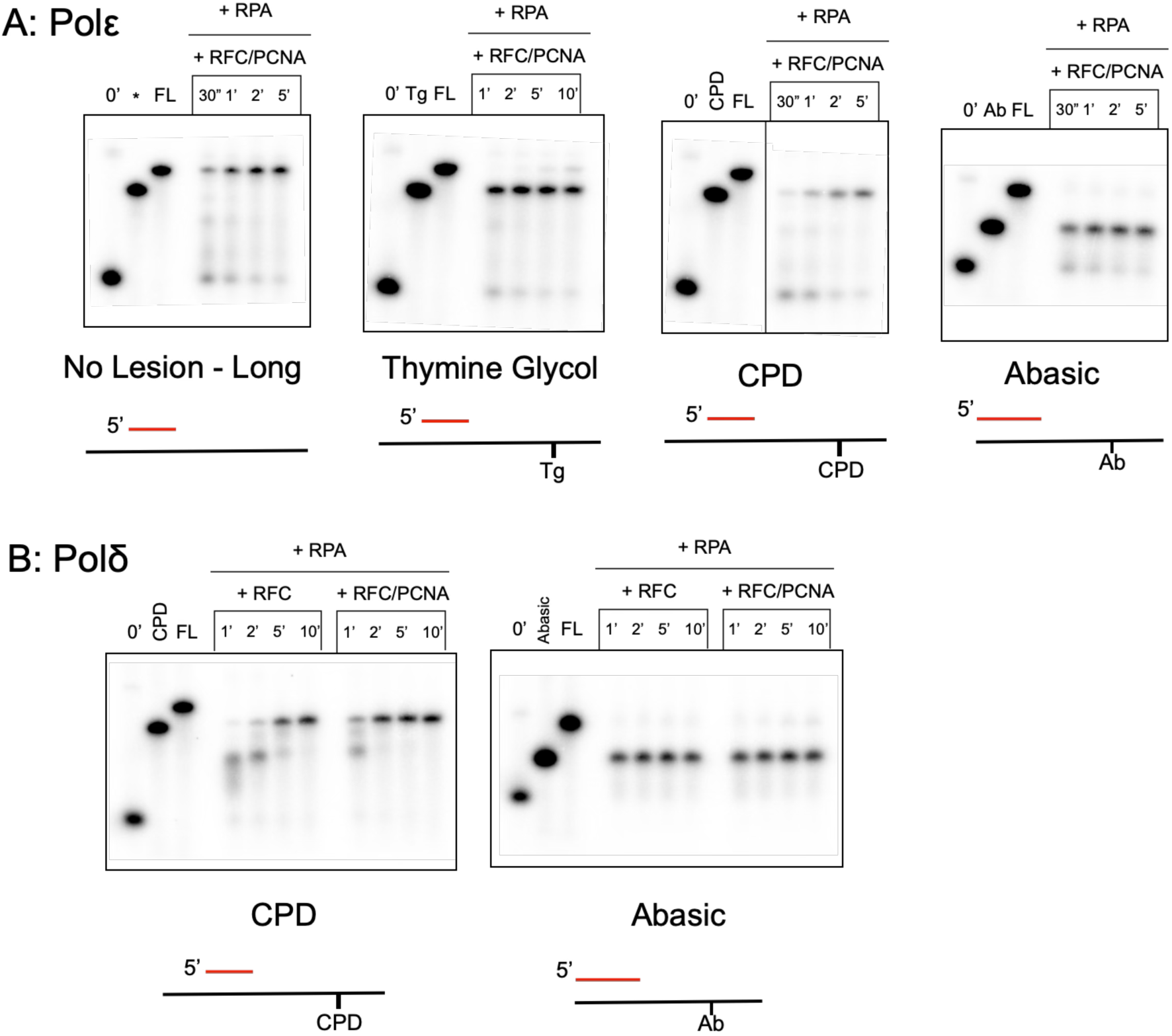
Impact of RFC +/- PCNA on extension and lesion bypass. Extension products are shown for assay with Polε or Polδ on the indicated substrate. In all cases, RPA was first loaded followed by initiation of extension by addition of polymerase, RFC, PCNA, ATP, and dNTPs. The * lane in no lesion substrate experiments indicates the size of products where stalling at a lesion would occur in the lesion carrying substrates. Red primers are ^32^P-ATP end-labeled.

**Figure S4:**
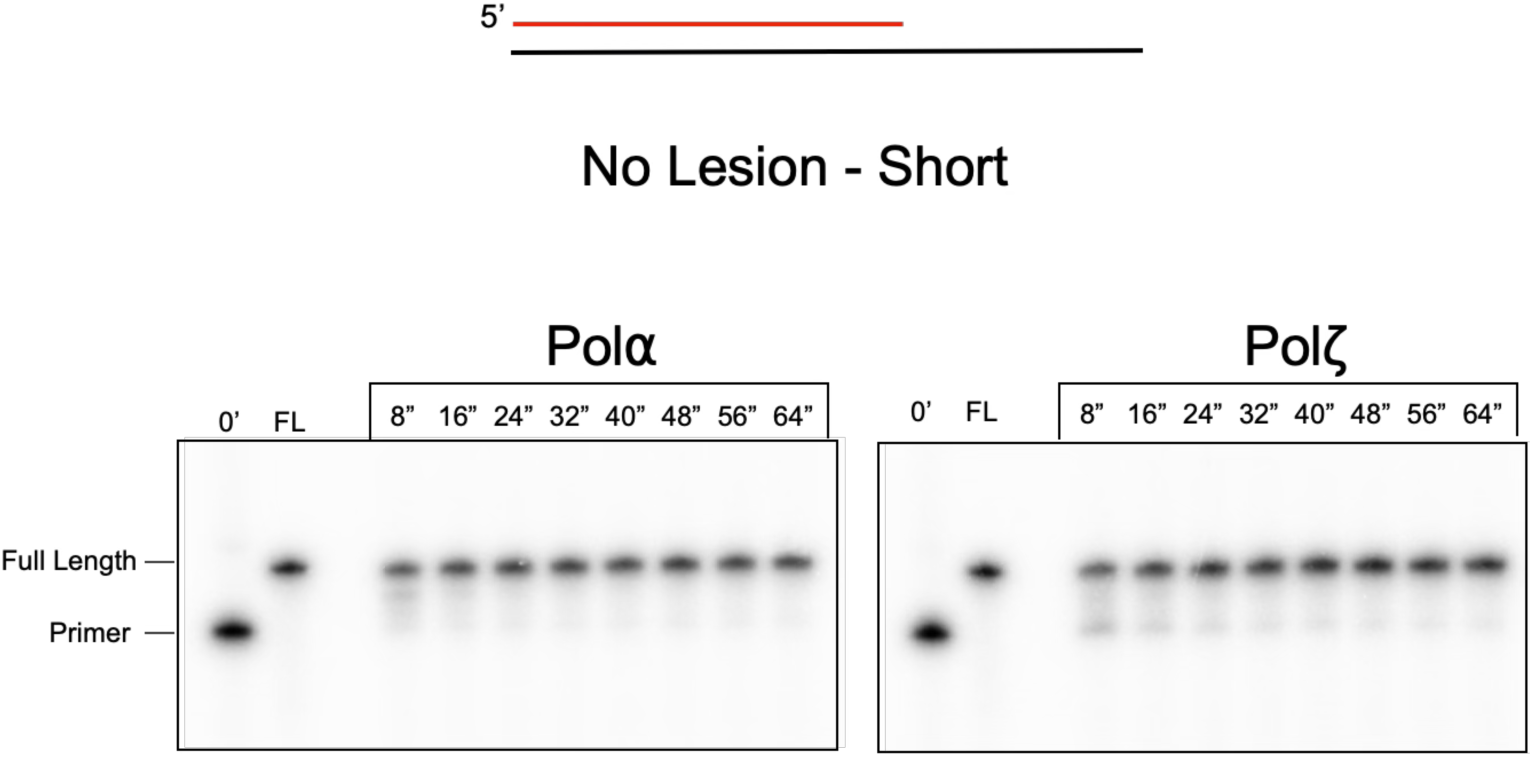
Primer extension on the short no lesion control substrate. Extension products from assays with Pol⍺ or Polζ at indicated timepoints. Unlike previous experiments, these are primed one nucleotide upstream of the site of lesions in parallel substrates. This illustrates that the 28 nucleotides of extension is completed very quickly. The red primer strand is ^32^P-ATP end-labeled.

**Table S1:**
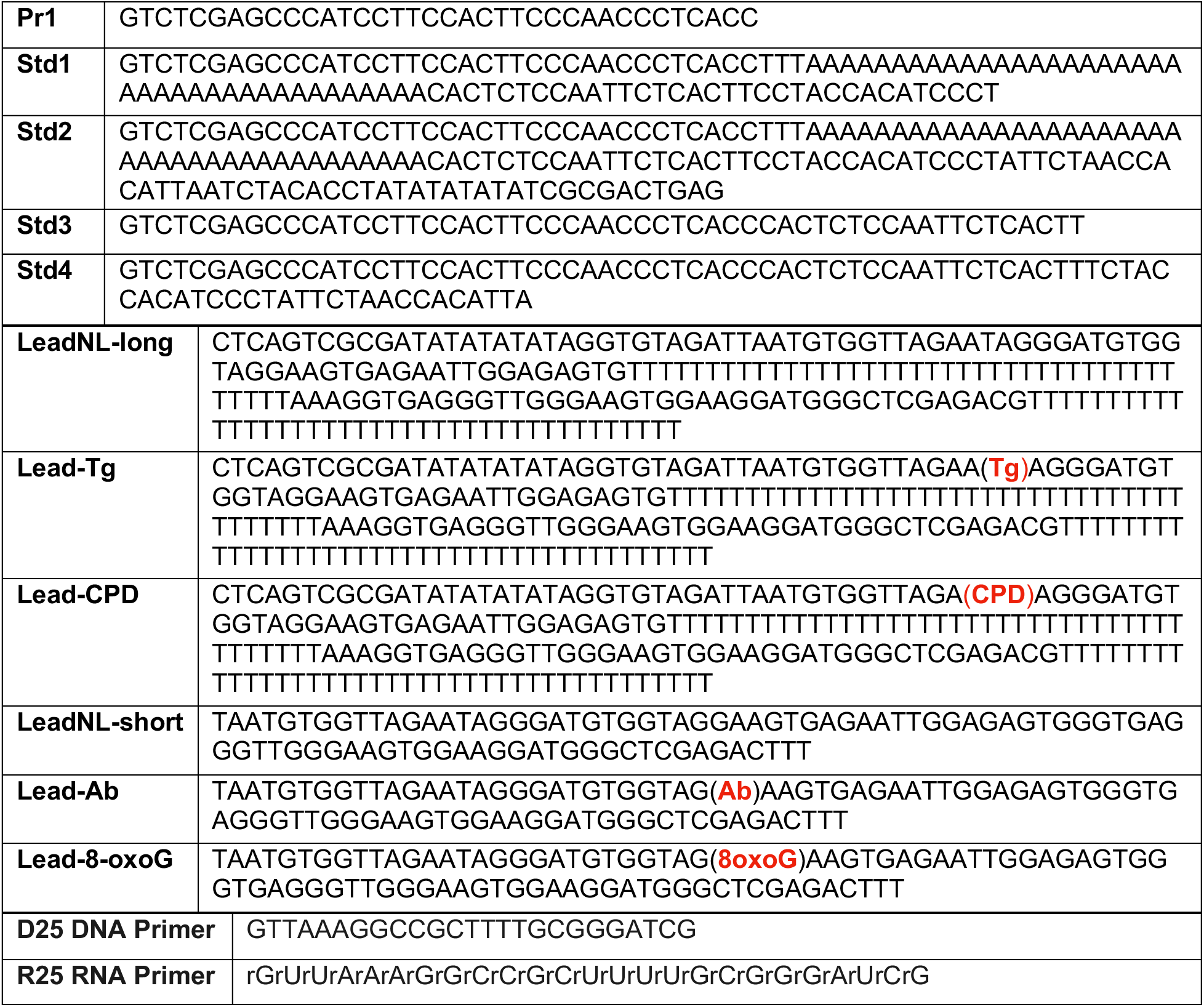
Sequences (5’=>3’) of oligonucleotides used in this study

